# Oxidation-reduction imaging of myoglobin unveils two-phase oxidation in the reperfused myocardium

**DOI:** 10.1101/2023.05.26.542538

**Authors:** Sally Badawi, Clémence Leboullenger, Matthieu Chourrout, Yves Gouriou, Alexandre Paccalet, Bruno Pillot, Lionel Augeul, Radu Bolbos, Antonino Bongiovani, Nathan Mewton, Thomas Bochaton, Michel Ovize, Meryem Tardivel, Mazen Kurdi, Emmanuelle Canet-Soulas, Claire Crola Da Silva, Gabriel Bidaux

## Abstract

Myocardial infarction (MI) is a serious cardiovascular problem that causes myocardial injury due to blood flow obstruction to a specific myocardial area. Under ischemic-reperfusion settings, a burst of reactive oxygen species is generated, leading to redox imbalance that could be attributed to several molecules, including myoglobin. Myoglobin is dynamic and exhibits various oxidation-reduction states that have been a subject of attention in the food industry, specifically for meat consumers. However, rarely if ever, have the myoglobin optical properties been used to understand the pathology of MI. In the current study, we develop a novel imaging pipeline that integrates tissue clearing, confocal and light sheet fluorescence microscopy, combined with imaging analysis, and processing tools to investigate and characterize the oxidation-reduction states of myoglobin in the ischemic area of the myocardium post-MI. Using spectral imaging, we have characterized the endogenous fluorescence of the myocardium and demonstrated that it aligns with the spectral profile of myoglobin. Under ischemia-reperfusion experimental settings, we report that the infarcted myocardium spectral signature is similar to that of oxidized myoglobin signal that peaks 3 hours post-reperfusion and decreases with cardioprotection. These results were correlated with MI measurements by Late Gadolinium Enhancement MRI. In conclusion, this seminal work suggests that the redox state of myoglobin can be used as a promising imaging biomarker for characterizing and estimating the size of the MI during early phases of reperfusion.

## INTRODUCTION

Cardiovascular diseases (CVDs) represent a major health and economic burden globally. Myocardial infarction (MI) is one of these diseases, characterized by severe myocardial injury resulting from partial or complete blockage of blood flow to a specific area in the myocardium, known as the area at risk. Restoration of this blood flow significantly influences the ischemic injury and leads to the progression of myocardial necrosis. This progression occurs in a wavefront phenomenon, moving from the subendocardium to the epicardium ^1–4^. The imbalance of oxygen supply during the myocardial ischemia-reperfusion process results in a massive amount of reactive oxygen species (ROS). ROS exacerbate myocardial injury by oxidizing cellular components^5^ and lead to redox imbalance in the heart.

Myoglobin (Mb) is a significant source for generating ROS during myocardial ischemia-reperfusion settings. Mb is expressed in high concentrations in the heart and the skeletal muscles (around 300µM and 4-5mM in terrestrial and marine mammals, respectively) ^6^. At the same time, it is present at much lower concentrations in smooth muscle. Like hemoglobin, Mb binds to oxygen molecules through a heme group and produces the OxyMb form. The latter is believed to function as a short-term reservoir of O_2_ ^7^ and/or to have a role in dioxygen transport to the mitochondria under hypoxic conditions ^6^. Mb can also bind to CO_2_ (CarbMb), facilitating its transport in exchange with O_2_. Moreover, Mb exhibits a nitrite reductase activity, which helps scavenge excessive nitric oxide (NO) and protects the cell from deleterious oxidation ^8,9^. During ROS-mediated oxidation of OxyMb, Mb is consequently oxidized, forming MetMb. Thus, as an allosteric enzyme, Mb can either be reduced or oxidized depending on the concentration of its various substrates.

Historically, Mb has been the subject of extensive research by the food industry due to its attractive role in pigmenting meat. The different redox states of myoglobin are responsible for the change in color of fresh meat from red to brown when exposed to light or high temperatures, which is a significant concern for meat consumers ^10^. Furthermore, the absorption spectra of myoglobin are highly sensitive and significantly affected by its interchangeable structure, which has been considerably studied to quantify proportions of the various forms and assess the quality of the consumed meat ^10–13^. Overall, Mb could thus potentially serve as a reporter to investigate the oxidation-reduction levels in cardiac and muscle cells.

Fluorescence imaging of unlabeled tissue is challenging and requires knowledge about endogenous fluorescent emitters. Prior studies reported that cleared mouse hearts displayed an amber-like appearance when illuminated with white light, which can be attributed to the presence of endogenous “pigments” or molecules that serve as “auto” fluorescent emitters ^14,15^. The total fluorescence intensity of a fluorophore is equal to the product of its molecular brightness and its concentration. Conversely, endogenous “auto” fluorescent emitters have low molecular brightness compared to fluorophores. As a result, endogenous “auto” fluorescence, in the absence of fluorochromes, mainly depends on the concentration of endogenous fluorescent emitter. Mb has been identified in cardiomyocytes as the most concentrated porphyrin, with an estimated concentration of around 300µM ^16^. In addition, Garry et al. showed that the pigments were strongly attenuated in the heart of Mb knock-out mice ^17^. At the cellular level, Mb localization spans from the cytoplasm, where it carries dioxygen ^18^, to mitochondria, where it acts as a buffer and reservoir of dioxygen ^19^. This suggested that myoglobin could significantly affect the myocardium’s endogenous fluorescence.

Optical tomography has demonstrated its ability to reconstruct the shape and macroscopic structure of the mouse heart in three-dimensions. It can also be combined with MRI images to enhance the image rendering of the infarct area ^20,21^. Multi-modal imaging, combining MRI, SPECT, and light sheet microscopy, has been proposed, but it has yet to display any significant improvement ^22^, where optical constraints limited all these historical studies. In this regard, optical properties of Mb have been utilized to evaluate its saturation through fluorescent excitation at 415 and 450 nm wavelengths in perfused rat hearts ^23^.

Furthermore, spectral deconvolution approaches utilizing Mb absorbance characteristics have been employed to evaluate oxygenation and mitochondrial redox in rabbit-perfused hearts ^24^. However, these studies were limited by inconsistent heart transparency, which was required for accurate quantification. In recent years, this limitation has been overcome by clarification techniques that render biological samples transparent and easily accessible for immunolabeling and three-dimensional volumetric imaging. Several clearing protocols have been applied to the heart for various purposes, including visualizing the myocardial vasculature ^25^ and examining the organization of fibers in the myocardial wall ^26^. Interestingly, in MI-induced model, tissue-clearing approach has been used to study the architecture of cardiac cells in the ischemic area ^27^ and investigate the phenotypic changes in cells, such as the fibroblasts, in the myocardium post-MI ^28^. An additional advancement was made when Merz et al. combined optical tomography for organ reconstruction based on its endogenous fluorescence with a fluorescent probe that diffused and labeled the injured area ^29^.

Little attention has been given to Mb’s endogenous fluorescence in this context. The latter’s signal can provide valuable information about the oxidation-reduction state of the myocardium. By measuring the fluorescence intensity of Mb in different oxidation states, one might determine the redox balance in the tissue and potentially use this information as a biomarker for disease or injury. However, the need for automated quantitative workflows is a challenge that must be addressed to fully exploit the potential of endogenous fluorescence imaging.

Our current study presents a novel automated imaging approach that integrates confocal and light sheet fluorescence microscopy, tissue clarification, and imaging tools to investigate and characterize myoglobin’s oxidation-reduction state in the myocardium’s ischemic area post-MI. To achieve this, we utilize the spectral characteristics of myoglobin and quantify the distribution of intensities in three-dimensional volumetric images at different time points following reperfusion. In addition, we evaluated mouse hearts protected by ischemic post-conditioning to demonstrate the specificity of our analysis in detecting changes in tissue oxidation.

## METHODS

### DATA AVAILABILITY

The scripts described in the Image Analysis section of Supplemental Methods are available on a public GitHub repository: https://github.com/chourroutm/Amira-Avizo_custom_modules).

Detailed methods and experimental procedures can be found in **Supplemental material**.

## RESULTS

### Myocardial endogenous fluorescence

The endogenous fluorescence properties of the myocardium can be preserved after clarification with aqueous-based and hydrogel embedding solutions. We developed and evaluated a new methodological pipeline, as depicted in **Figure 1A**, to determine whether the endogenous fluorescence of the heart could serve as a reliable indicator to measure the degree of infarct injury resulting from ischemia-reperfusion. A mouse model of MI was induced by occluding the left-anterior coronary artery for 1h, followed by reperfusion for different periods (15min, 3h, and 24h) until the mice were sacrificed ^30^ (**Supplemental Figure S1**). The mice included in the study were categorized into three groups: a negative control group composed of 24h sham mice, an ischemic group at different time post-reperfusion, and a positive control group for cardioprotection represented by mice undergone ischemia-reperfusion for 24h with an ischemic post-conditioning (IPoC) at the onset of reperfusion. The hearts of the mice were collected, fixed, and processed for standard X-clarity protocol without bleaching by hydrogen peroxide. Details of the process are provided in the supplemental methods section. Hearts were then imaged using either confocal microscopy or light sheet microscopy, and the resulting images were then processed and analyzed using various software. It is worth noting that while the clearing procedure did not achieve complete transparency of the myocardium, it significantly increased the depth of photon penetration (**Figure 1B**). By utilizing multiple lasers with excitation wavelengths of 405 nm, 488 nm, 561 nm, and 633 nm and capturing the resulting light with a spectral detector, specifically a single photon avalanche photodiode (SPAD), a wide fluorescence bandwidth was detected (**Figure 1C**). This endogenous fluorescence of the myocardium provided high-quality macroscopic imaging of myocardium structures and myofibrils (**Figure 1D**).

**FIGURE 1.**
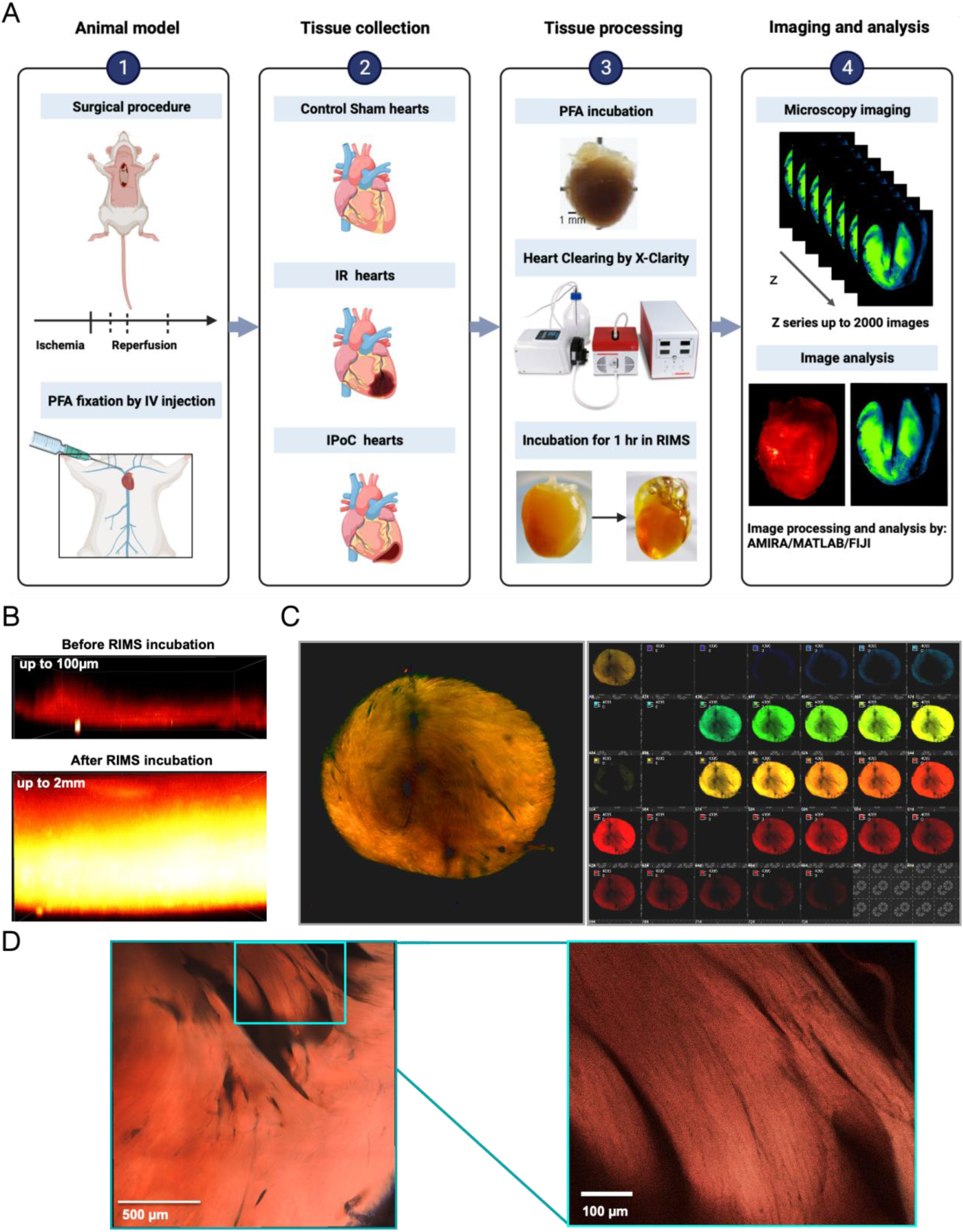
Cleared mouse hearts conserved endogenous fluorescent emitters in the absence of hydrogen peroxide bleaching. **A.** Methodological pipeline of the study: a mouse model of myocardial infarction, mouse heart before clarification (top), after clarification with the X-Clarity system (bottom right) and after incubation of the cleared heart in the mounting medium (bottom left), fluorescence imaging of unlabeled hearts and image analysis. **B.** A 10-fold increase in photon penetration in the depth of the myocardium could be measured in the cleared heart by a mono-photonic confocal microscope under excitation at 488nm. **C.** A 16-tile image of a cleared myocardium excited by 4 laser lines: 405/488/561/633nm and acquired with a spectral detector showed endogenous fluorescence over a large wavelength band. Split image (right) and a composite image of merging the 32 channels are shown to the left. **D.** Tiles image of a spectral acquisition acquired tangentially endocardium side of the left ventricle shows the papillary muscles. Inset: magnification of the papillary muscles enables the identification of single fibers.

### Structure of the endogenous fluorescence signal in cardiomyocytes resembles myoglobin localization

By focusing on the red wavelength range (bandpass >620 nm), expected to be partly generated by myoglobin fluorescence, and optimizing the resolution using a confocal microscope, we revealed various myocardial structures such as artery layers, cardiomyocytes and interstitial cells (**Figure 2A**). In addition to the cellular phenotypes, the endogenous fluorescent signal exhibited a distinct grid-like pattern within the cardiomyocytes (**Figure 2B**). This pattern is typically associated with the ultrastructure of cardiomyocytes, including t-tubules, sarcoplasmic reticulum, mitochondria, and myofilaments. We then aimed to investigate if Mb could be responsible for generating this spatial distribution. We have immunostained isolated cardiomyocytes with Mb and GRIM-19, a mitochondrial protein marker (**Figure 2C**). Overlay of Mb and GRIM-19 signals revealed that Mb colocalized with GRIM-19 at the periphery of mitochondria and spread away from mitochondria in a well-ordered grid-like pattern resembling the organization of the t-tubules (**Figure 2D**). Overall, immuno-detection of Mb exhibited a spatial signature that closely resembled the endogenous fluorescence detected in cleared hearts’ cardiomyocytes. Taken together, our results and previous research support the hypothesis that Mb could be one of the primary sources of endogenous fluorescence in cardiomyocytes, specifically in the red wavelength range.

**FIGURE 2.**
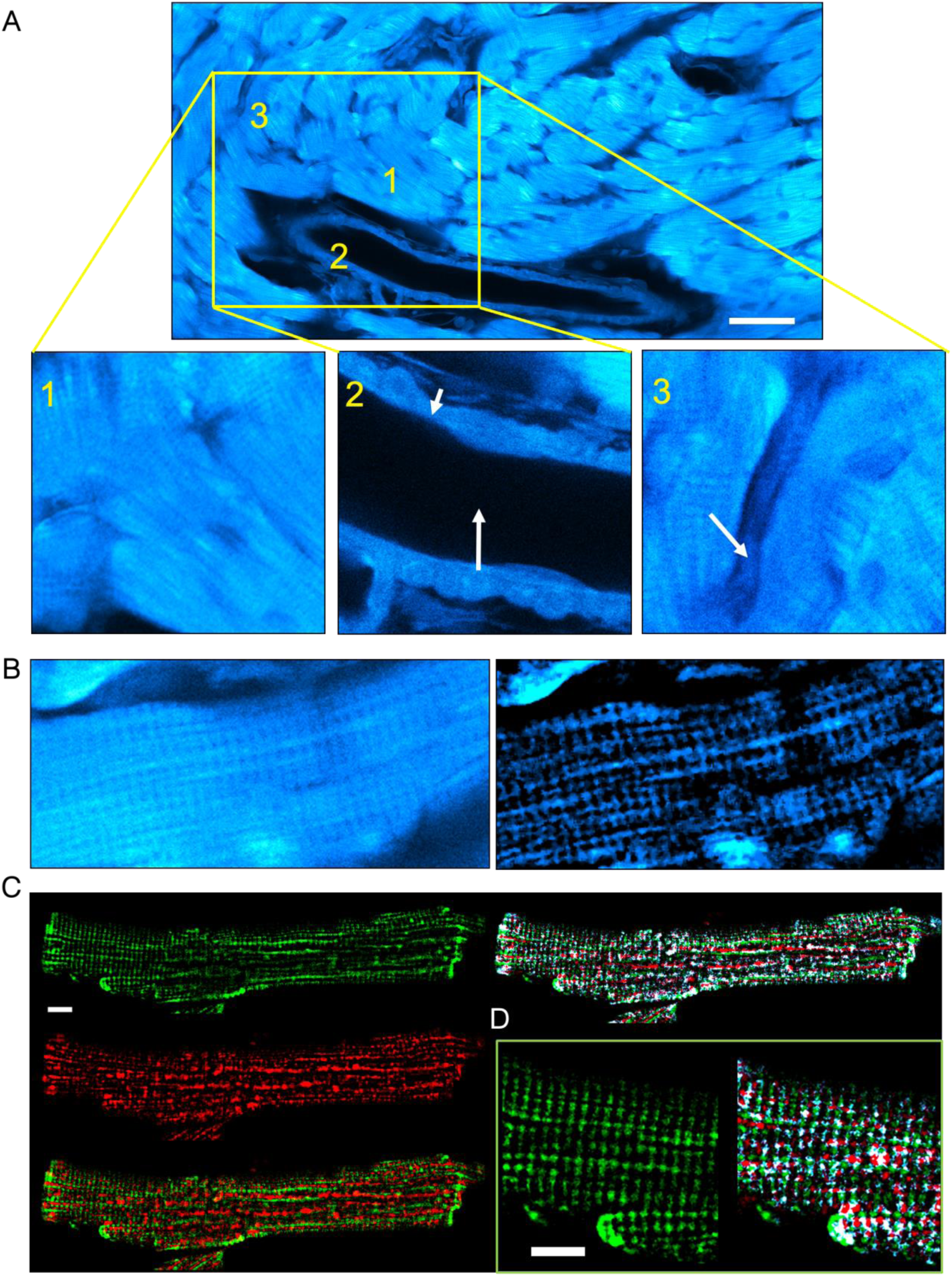
Myoglobin localization in cardiomyocytes correlates with the subcellular structures emitting endogenous fluorescence in cleared mouse hearts. **A.** Images of a cleared unlabeled mouse heart taken with a confocal microscope (60x objective, laser source at 633nm, and SPAD detector) at maximal resolution. Insets 1, 2, and 3 show magnification of cardiomyocytes, aorta, and an interstitial cell. Scale bar: 50 µm **B.** Higher magnification of a single cardiomyocyte section before (left) and after (right) entropic filtering. **C.** Images of adult mouse cardiomyocyte immunolabelled with anti-myoglobin (green channel) and GRIM-19 (red channel). The overlay is shown on the right, and the pixels with both fluorescence signals are white. **D.** Magnification of the image shown in C.

### The oxidized myoglobin’s spectral signature recapitulates the infarcted area’s spectral signature in failing hearts

Peroxynitrite or hydrogen peroxide can efficiently oxidize the iron atom in the heme group of Mb, forming oxidized myoglobin (MetMb). Over-oxidation of MetMb leads to the formation of ferryl myoglobin (ferryl-Mb), whose absorption spectrum is similar to that of MetMb. We utilized the spectral detector of a confocal microscope to assess how the changes in its oxidation-reduction state modified the fluorescence emission spectrum of pure horse myoglobin. Initially, we had to eliminate the gaps of the dichroic mirrors 405/488/565/647 nm and the glass reflection to prevent the bleed-through phenomenon observed at the peripheries of the 647 nm dichroic mirror (**Supplemental Figure S2A**). Next, we conducted a dose-effect and time-effect study using hydrogen peroxide (H_2_O_2_) to determine the optimal dose required for MetMb ^31^. As displayed in **Supplemental Figure S2B**, a dose of 0.03% H_2_O_2_ was deemed appropriate for oxidizing Mb without inducing any signal bleaching or destruction. As demonstrated in **Figure 3A**, enrichment in reduced myoglobin (CarbMb) decreased fluorescence emission intensity, whereas enrichment in MetMb increased it compared to oxygenated myoglobin (OxyMb). Assuming that Mb indeed contributed to the endogenous fluorescence of myocardium, we postulated that alterations in the oxidoreduction state of myocardium would exhibit a comparable signature in myoglobin’s spectra. Pretreatment of control hearts with H_2_O_2_, without clearing, revealed a significant increase in the endogenous fluorescence intensity above 600 nm (**Figure 3B).** Interestingly, using sodium dithionite to reduce Mb prior to clearing slightly increased endogenous fluorescence intensity around 650 nm, with no effect in greater wavelengths, unlike H_2_O_2_. Notably, the observed fluorescent signal around 650 nm was suppressed after heart clearing, suggesting it was likely due to lipids (**Figure 3C)**. The mean oxidation of myocardium was associated with an increase in endogenous fluorescence intensity above 600 nm, while reduction by sodium dithionite had no significant effect or slightly decreased it. Additional controls assessing treatments’ time and concentration are provided in **Supplemental Figures S2C and S2D**. In addition, spectral profiles of the fluorescence of individual hearts are given as examples in **Supplemental Figure S2E**. Altogether, these findings indicate that myoglobin’s oxidation can partly explain the shift in the endogenous fluorescence spectrum and will be referred to as the spectral signature of MetMb.

**FIGURE 3.**
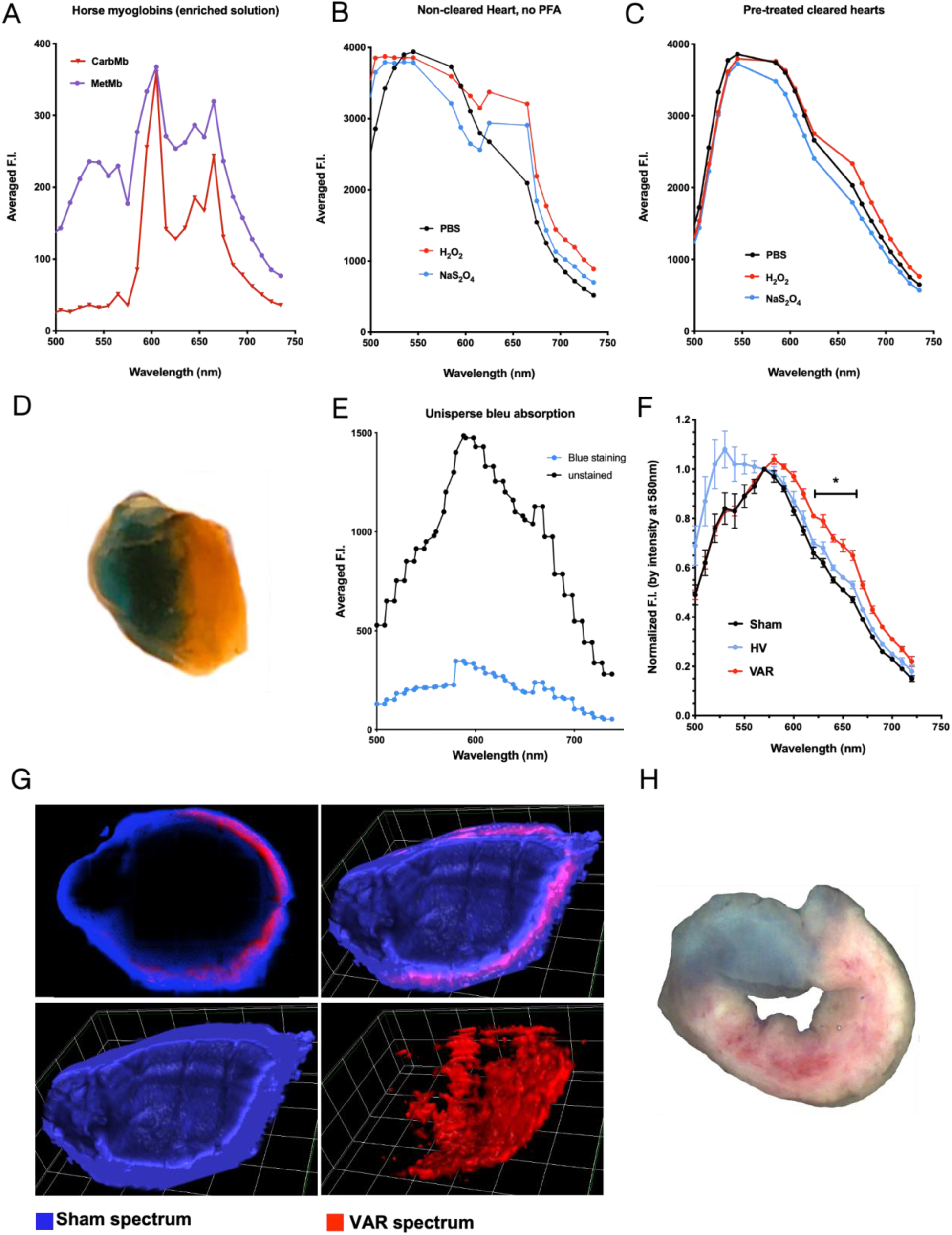
Myoglobin oxidation and myocardial ischemia-reperfusion share a spectral shift in the red wavelength range. **A.** Fluorescence spectra of pure horse myoglobin measured with the confocal microscope under illumination with laser lines: 405, 488, 561, and 633nm and detected by a spectral avalanche photodiode. Myoglobin was enriched in either the form bound to CO_2_ (CarbMg), or the oxidized form (MetMg). **B.** Fluorescence spectra of uncleared heart incubated either in PBS, hydrogen peroxide to induce oxidation (H_2_O_2_), or dithionite to induce reduction (NaS_2_O_4_) were determined as in A. **C.** Fluorescence spectra of heart incubated either in PBS, H_2_O_2_ (oxidation) or NaS_2_O_4_ (reduction) before clarification were determined as in A. **D.** Image shows a cleared I/R mouse heart labeled with Unisperse blue (healthy area) and contrasting with the brownish area-at-risk. **E.** Fluorescence spectra of healthy area (labeled by Unisperse blue) and area-at-risk determined as in A. **F.** Normalized fluorescence spectra of healthy volume (HV, blue line) and volume-at-risk (VAR, red line) of a 24h-reperfused heart and a sham heart (black). Value: Mean±SD; n=3 Sham and 4 IR. * p<0.05. **G.** Images of a 24h-reperfused heart by linear unmixing of the mean spectra of sham and VAR presented in F. Confocal slice (top left), 3D rendering volume of healthy volume (bottom left), volume-at-risk (bottom right) and combined volumes (top right). **H.** Heart slice labeled by Unisperse blue (healthy area) shows the distribution of dead tissue (white) and living tissue (pink) in the area-at-risk as labeled by a TTC assay.

Next, we then took advantage of Unisperse blue labeling (**Figures 3D and 3E)** to extract endogenous fluorescence spectra from both healthy myocardial volume and the whole occluded myocardial volume, defined as “volume-at-risk” (VAR). The control spectrum was retrieved from a sham heart without Unisperse blue. Following normalization by the peak value of fluorescence intensity (at 585 nm), the mean endogenous fluorescence spectrum in the VAR exhibited a significant increase of fluorescence intensity above 600 nm, with a maximal difference from Sham and healthy volume around 650 nm (**Figure 3F).** Except for the specific signature of Unisperse blue in the 500 nm bandpass, Sham and Healthy hearts showed similar endogenous fluorescence spectra. This increased fluorescence intensity in the bandpass 620-660nm in the VAR resembles the shift of in vitro spectral signatures between CarbMb and MetMb. While emission spectrum analysis of images is a powerful technique, it is subjected to significant drawbacks when light passes through a diffractive or absorbing media: a red shift in the light spectrum is anticipated to occur at greater depths due to the lower penetrance of the shorter wavelengths. In the cleared heart samples, we were able to control for this effect and detected only a slight occurrence on the inner border of the myocardial wall (**Supplemental Figure S3A**).

We then applied spectral linear unmixing to segment sham-like spectrum and VAR spectrum in all hearts (**Supplemental Figure S3B)**. The spectral signature of VAR, which resembles the spectral signature of MetMb, was detected at the expected position of VAR in the left ventricle. However, it did not propagate transmurally but radiated from the middle of the myocardium wall (**Figure 3G)**, what is similar to the necrotic area (white area) detected in the TTC assay (**Figure 3H**). This finding excludes the possibility that the red shift phenomenon caused the detected signal due to in-depth photon absorption. In addition, this suggests that the fluorescence spectrum in the VAR is enriched in MetMb spectrum. By contrast, some random noise signal was detected in sham hearts (**Supplemental Figure S3C**).

Altogether, our results suggest that the spectral features and spatial distribution of the endogenous fluorescence above 600 nm in infarcted mouse hearts fit with myoglobin’s oxidoreduction state and location within the infarcted volume. Therefore, we next aimed to establish a pipeline to quantify the oxidized volume characterized by MetMb fluorescent signature in hearts collected at different reperfusion times and the decrease in the oxidized volume induced by ischemic post-conditioning (IPOC).

### The intensity of endogenous fluorescence at 633 nm is associated with reperfusion time, and cardioprotection by IPoC, with a stronger oxidation signal observed 3 hours post-reperfusion

In comparison to the z-stacks generated by confocal microscopy, the spatial resolution and imaging depth have been drastically improved using light sheet microscopy (**Supplemental Figure S4A**), particularly in the Z plane, with a difference of 100µm versus 4µm, for image files of similar size (50-100Go), with the notion that spectral emission was not possible. All the hearts were imaged using light sheet microscopy at a wavelength of 633 nm, and the light was collected with an extended pass filter above 650 nm. The first step of the analysis involved anatomical segmentation of the left ventricle. Subsequently, using the strong contrasting effect of the Unisperse blue in the healthy area, we extracted the volume-at-risk (VAR) with intensity-based thresholding (**Figure 4A)**. Voxels from VAR were then plotted on a distribution histogram (**Figure 4B),** which revealed apparent differences between the experimental groups. This difference in intensity distribution among the voxels was accompanied by variations in the spatial organization of the oxidation signal (**Figure 4C)**. However, robust and objective signal quantification would be required to demonstrate significant and reproducible differences in the oxidized signal between experimental conditions.

**FIGURE 4.**
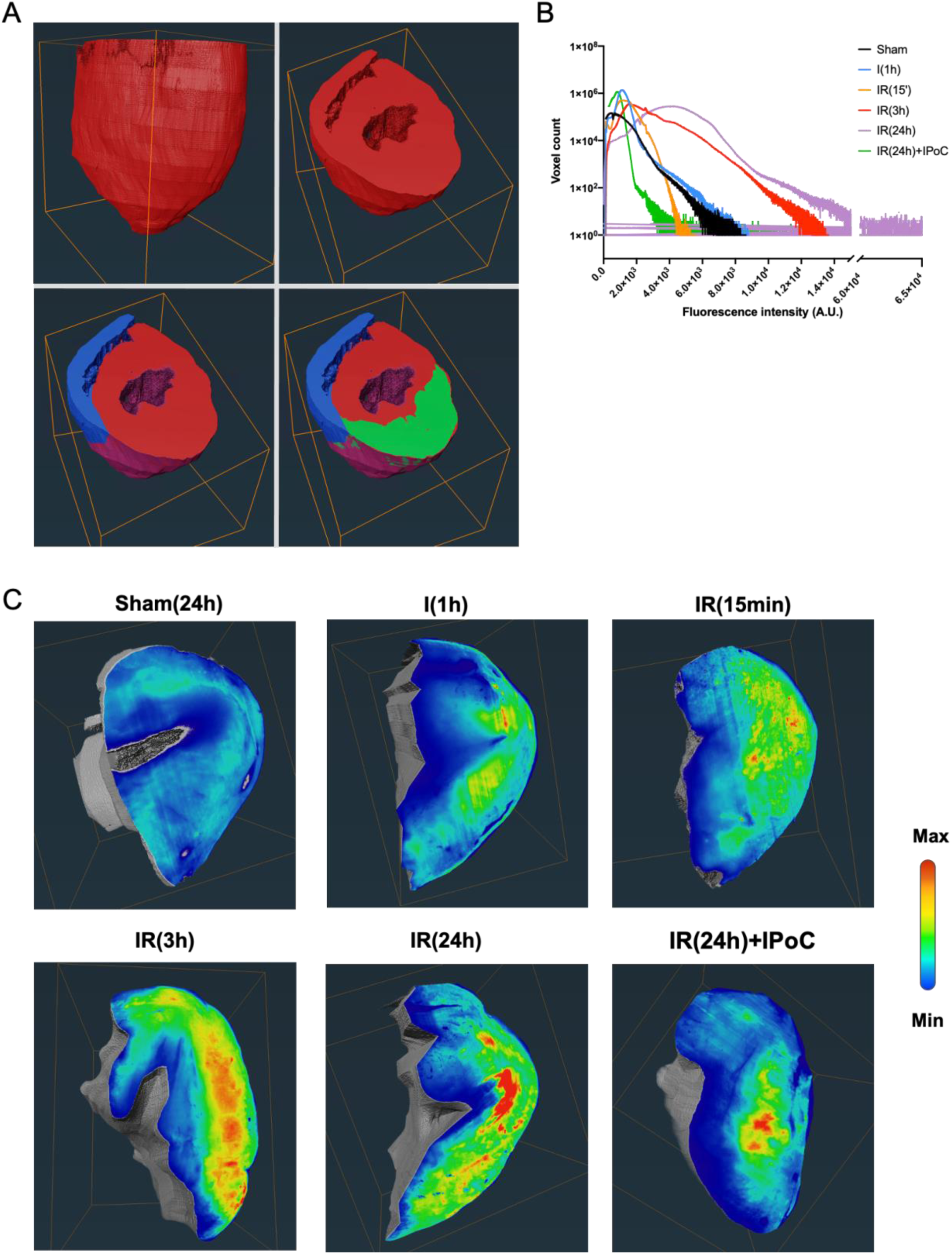
Light sheet imaging at 633nm reveals hypersignal endogenous fluorescence in the area-at-risk. **A.** Stack of images obtained by an ultramicroscope at 633nm were segmented as represented: the right ventricle (blue) left ventricle (red) and volume-at-risk (VAR). All hearts excepted the sham were labeled by Unisperse blue, which was used to retrieve the VAR. An equivalent section was selected in the sham hearts to compare fluorescence intensity on a similar volume. **B.** Distribution histograms of VAR voxels by fluorescence intensity (1 VAR per histogram). **C.** Medial transversal slice of the 3D segmented VAR were colored such as deep blue labeled the minimal fluorescence intensity value of the histogram and red labeled the maximal fluorescence intensity. A representative heart is shown for each group: sham heart (no infarct) 24h after the surgery: sham(24h), heart after 1h ischemia: I(1h)), reperfused heart at 15 min, 3h and 24h: IR(15min), IR(3h) and IR(24h), respectively; and a heart subjected to ischemic post-conditioning and reperfused for 24h: IR(24h)+IPoC.

There were two major concerns that needed to be addressed in this analysis: 1) variability in the degree of clarity among the hearts, resulting in heterogeneity in the photon absorption along the light spectrum, and 2) heterogeneity in the light path due to differences in myocardium wall length, ventricular lumen, and, excepted in sham hearts, the presence of Unisperse blue in the healthy area (**Supplemental Figure S4B)**. These factors contributed to experimental noise that made it challenging to use an intensity-based segmentation to analyze the oxidation signal accurately. To overcome this, we aimed to develop a more robust analysis flow to compare the intensity distribution of the oxidized signal, defined an objective segmentation threshold and finally quantify the oxidized volume in all hearts, individually. Sham hearts were not included in the first phase because of the lack in Unisperse blue which slightly modifies the distribution of the fluorescence intensity as compared to the end of the ischemic phase (**Figure 4B**). To normalize the experimental noise, we deleted low intensities (background noise), removed low-frequency intensities at the foot of the distribution at a fixed threshold of 0.002%, and normalized intensity values between 0 and 1 (where 0 is the lowest intensity voxel and 1 is the highest intensity voxel) (**Supplemental Figure S5A-D**). A frequency histogram was thus plotted by dividing each column value by the total number of voxels in VAR (**Supplemental Figure S5E**). These frequency histograms were used to normalize VARs before averaging. Frequency histograms of the same experimental groups were then averaged (**Supplemental Figure S5F**). Finally, after testing multiple Gaussians fits, we found that the model with two Gaussian best fits the mean frequency histograms (**Supplemental Figure S5G and Figure 5A**). As demonstrated in **Table 1**, the increase in higher fluorescence intensities with reperfusion time was confirmed by a decrease in areas under the curves (AUC) of the left Gaussian fit (lowest fluorescence intensities), an increase in the AUC of the right Gaussian fit (highest fluorescence intensities), and a decrease in the overlap between the AUC of the two Gaussian fits. In the first section of our study, we demonstrate that the oxidation signal generated by MetMb lies in the increased intensity of endogenous fluorescence in the red wavelengths. As a result, we defined the right Gaussian fit as the best estimator of the shift from CarbMb to MetMb. We conducted a comparison among the different experimental groups. The median of the right Gaussian fit was 0.485 towards the end of the ischemic period; it rose to 0.527 after 15 minutes of reperfusion and increased to 0.566 after 3 hours of reperfusion. Interestingly, the median of the right Gaussian fit at 24h after reperfusion was similar to that obtained with IPoC (0.490 and 0.477, respectively). In summary, these results suggested that the oxidation intensity increased during reperfusion, reached its maximum by 3h of reperfusion, and then decreased. This observation correlates with the blood level of Troponin I used in clinical settings to estimate infarct size and evaluate reperfusion lesions (**Figure 5B**). The correlation over time between blood troponin level and MetMb fluorescence intensity was almost perfectly linear with different slopes from ischemia to 3h reperfusion and 3h to 24h reperfusion (**Figure 5C)**. Only two median values: IR(3h) and IR(24h) of MetMb fluorescence intensity were found above the mean of the median of all experimental groups (0.509) and only the condition IR(3h) had blood troponin level above the mean value of all experimental groups (37,985). Finally, in addition to measuring the intensity of oxidation, our approach aimed to quantify the oxidized volume.

**FIGURE 5.**
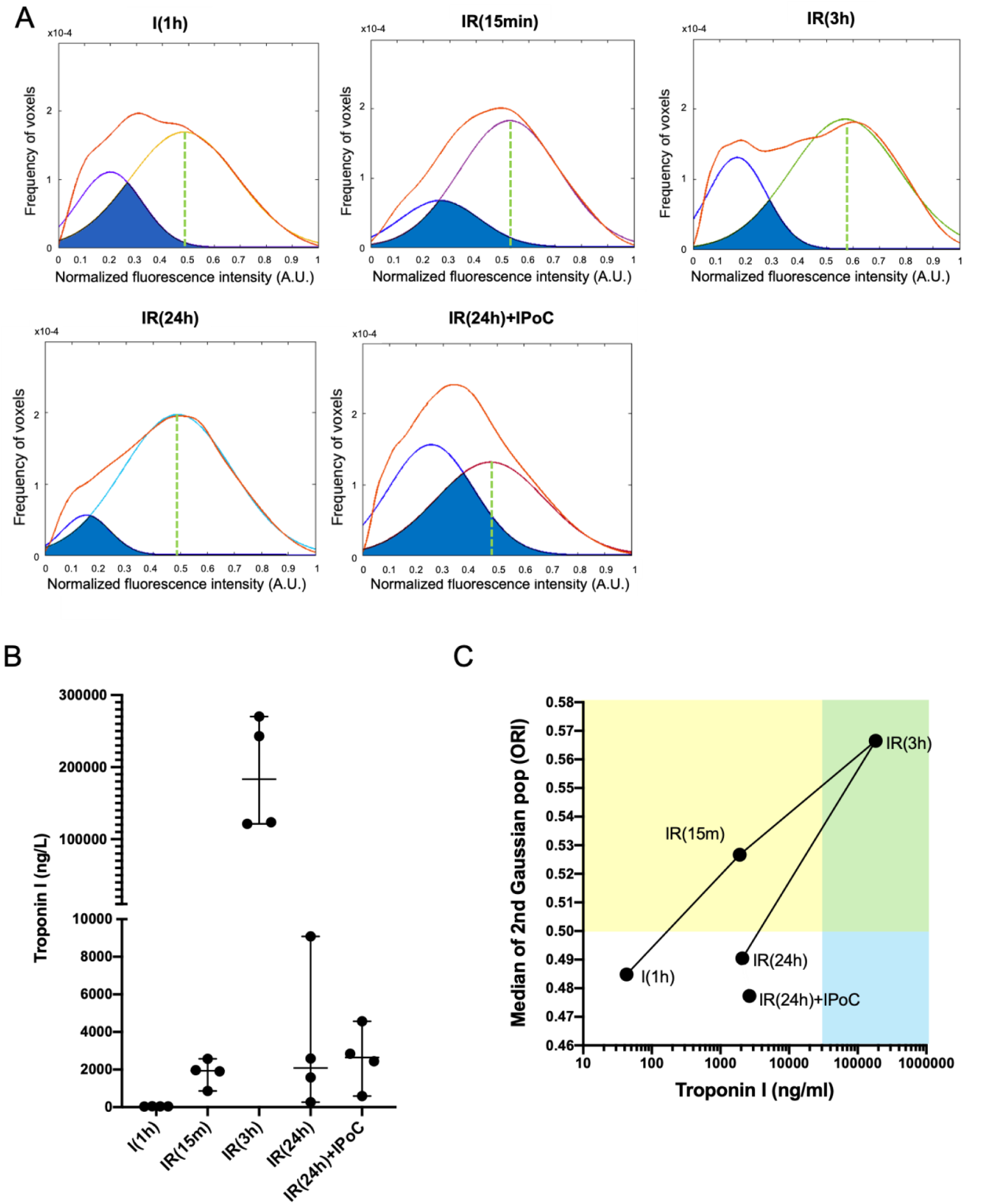
Quantification of oxidation intensity at reperfusion by oxidation-reduction imaging. **A.** Mean voxel distribution histogram of fluorescence intensity per experimental group (n=6 AAR per group) is shown in red. The two Gaussian fits of the mean distribution histogram: left Gaussian fit for the lowest fluorescence intensities and right Gaussian fit for the highest fluorescence intensities model the proportion of CarbMb and MetMb. The green dotted line shows the median of normalized right Gaussian fit. Fluorescence intensity **B.** Blood troponin I concentration assayed by ELISA (n=4 per group). **C.** Biplot showing median value of blood troponin concentration in function median value of normalized MetMb fluorescence intensity. Straight line connects time points of the pseudo-kinetic. The yellow area indicates MetMb fluorescence intensity above the mean of the medians of all experimental groups. The green area indicates blood troponin level above the mean of the medians of all experimental groups. The bleu area reports junction between yellow and green area.

**Table 1:** Gaussian fit values of the various experimental groups. Features of the Gaussian fits of the average frequency histograms and the two Gaussian population models for each experimental group.

| Group | Surface of 1<sup>st</sup><br>peak | Surface of 2<sup>nd</sup><br>peak | Surface of curves<br>Overlap | Mean residual between<br>fit and curve |
| --- | --- | --- | --- | --- |
| I(1h) | 0.27 | 0.73 | 0.19 | 0.40 |
| IR(15min) | 0.22 | 0.78 | 0.17 | 0.22 |
| IR(3h) | 0.27 | 0.74 | 0.10 | 0.82 |
| IR(24h) | 0.11 | 0.89 | 0.09 | 0.30 |
| IR(24h)+IPoC | 0.47 | 0.53 | 0.27 | 0.51 |

### The oxidized volume is maximal at 3h but is sustained up to 24h post-reperfusion and correlates with infarct volume measured by Late Gadolinium Enhancement MRI

As the median of right Gaussian fit at 1h ischemia, 24h reperfusion, and IPoC were similar and all below the mean of the medians, we used their averaged value (0.485) as a segmentation threshold above which the fluorescence intensity most probably come from MetMb molecules. This threshold value was used to segment the oxidized volume within each individual VAR, including those from sham hearts. The right Gaussian fit spanning from 0 to 1 in the normalized histograms, a threshold at 0.485 is equivalent to a threshold at 48.5% of the maximal intensity in intensity histograms. As shown in **Figure 6A**, segmentation of sham hearts revealed a patchy signal that may be either attributed to steady-state oxidation level in healthy myocardium or statistical errors in the image processing analysis. At the same time, segmentation of the oxidized volume in the VAR of a 24h-reperfused heart resulted in a transmural hypersignal consistent with the shape and localization of the necrotic area observed by TTC assay and shown in **Figure 3H**. It is noticeable that the patterns of the oxidized signal in sham and IR (24h) hearts, respectively, are similar to the ones detected by linear unmixing of myoglobin spectra in confocal images (**Supplemental Figure S3C**). The mean ratio of oxidized volume normalized by VAR shows a slight increase during ischemia and a significant increase 3h after reperfusion (**Figure 6B**). This value remained stable at 24h post reperfusion, supporting that although the peak intensity of oxidation was reached locally at 3h, the volume of tissue oxidized expanded in the first hours but was kept stable at least for 24h post-reperfusion.

**FIGURE 6.**
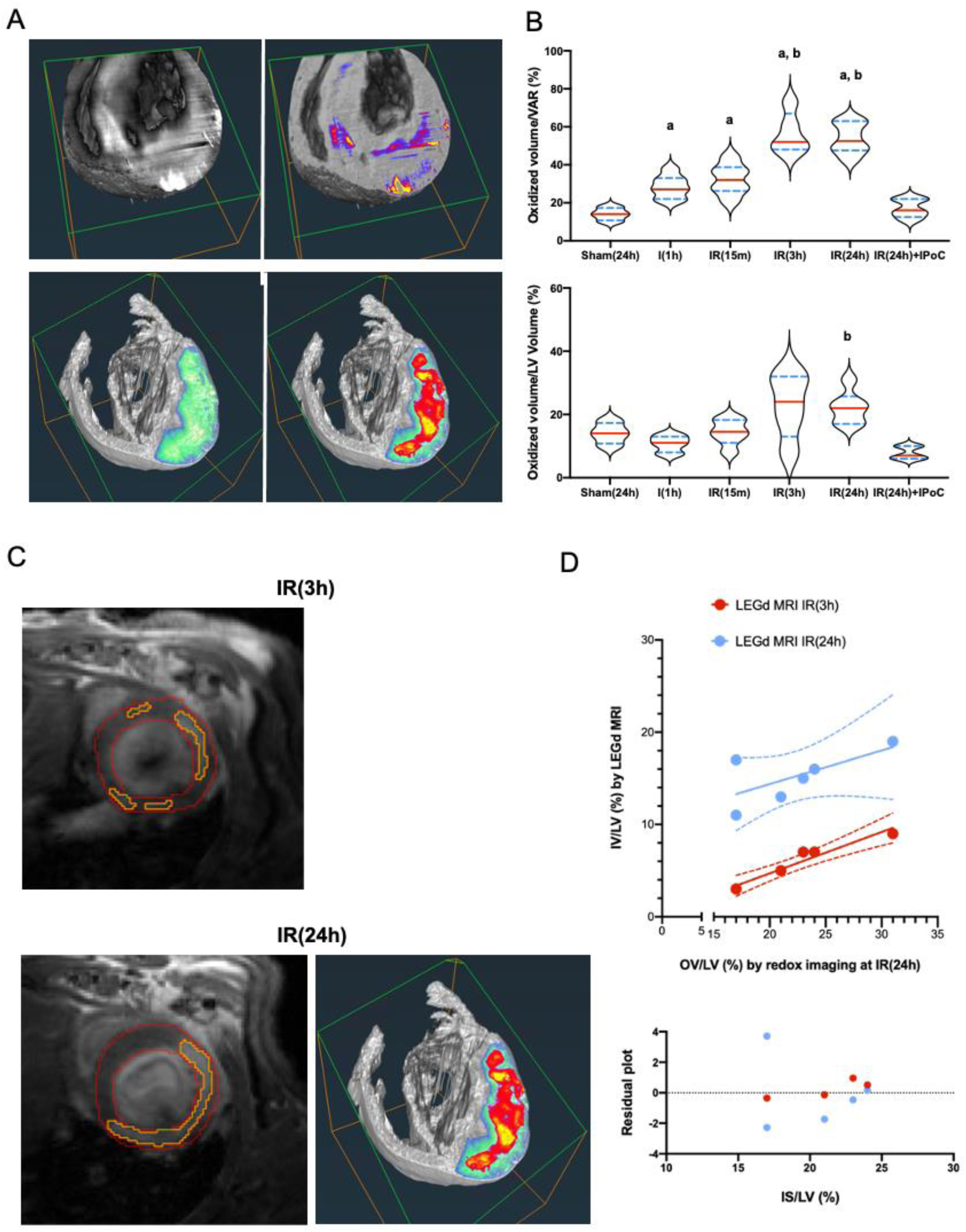
Quantification of the oxidized volume extension during reperfusion and its correlation with infarct size measure by MRI. **A.** 3D rendering volumes of sham (top panels) and 24h-reperfused heart (bellow). VAR is shown in a green-fire blue color scale; oxidized voxels are colored with the hot color scale. **B.** Plots represent values for the oxidized volume divided by VAR (top) or left ventricle volume (down) for each heart in each experimental group. Red line: median; blue dotted line: IC95%; n=6 per group. Significant difference with Sham(24h): a; Significant difference with I(1h): b was assessed by non-parametric Brown-Forsythe post-test and Welch ANOVA test. **C.** Images acquired by Late Gadolinium Enhancement MRI at 3h and 24h post-reperfusion in a living mouse and by oxidoreduction imaging post-mortem. **D.** (Top) Correlation plot showing the linear correlation between infarct size (at 3h and 24h post-reperfusion) and oxidized volume, normalized by left ventricle volume and (down) fit residual

Moreover, the emergence of this oxidized myocardial volume was prevented by IPoC. Noteworthy, the mean ratio of oxidized volume normalized by VAR at 3h and 24h post-reperfusion is consistent with the value measured by TTC in this mouse model. A ratio of oxidized volume normalized by the left ventricle volume was calculated to compare with values obtained from Late Gadolinium Enhancement MRI (**Figure 6B**). The Late Gadolinium Enhancement MRI was performed on the same animals subjected to late 3h and 24h reperfusion (**Figure 6C**). The values obtained from ORI and MRI were plotted on the same graph, displaying a good correlation irrespective of the reperfusion time (**Figure 6D**). However, while the maximal infarct volume measured by MRI was observed at 24h post-reperfusion, the maximal oxidized volume was already detectable at 3h post-reperfusion.

In summary, we demonstrated that the duration of reperfusion is associated with detectable oxidation of Mb in cleared and unlabeled mouse hearts, as determined by measuring the endogenous fluorescence through intensity or spectral imaging. We also established a reliable and robust analytical pipeline to quantify the intensity and volume of oxidized myocardium. We confirmed these values with blood levels of Troponin I and infarct volume determined by MRI.

## DISCUSSION

Myoglobin is released by injured myocytes in the heart and skeletal muscles. Plasma Mb levels were previously utilized to diagnose MI by measuring its release over time ^32,33^. However, this approach was abandoned in 2014 after the development of high sensitive Troponin I assay ^34^. Recently, a study suggested that plasma Mb levels could be used as a diagnostic marker of acute myocarditis ^35^. While few attempts have been made to image fluorescence signals in vivo, these studies have mainly focused on estimating tissue oxygenation levels in rat models of permanent ischemia ^36^ or skeletal muscles ^37^. Our study is the first to provide a multiscale imaging of Mb in the myocardium and to highlight its significance in the myocardial endogenous fluorescent signal. We also provide a seminal proof-of-concept that oxidation-reduction imaging (ORI) of Mb can be used to quantify the intensity of oxidation, the volume of oxidation, and the effect of cardioprotection in myocardial infarction.

### Fluorescence imaging of the unlabeled heart

Previous studies have reported that imaging of the macroscopic structure of the heart can be conducted through a fluorescence microscope, utilizing the endogenous fluorescence of the tissue ^29,38^. The origin of this endogenous fluorescence has been studied in various cells, tissues, and organisms using fluorescence spectroscopy (for review, see ^39^). Collagen, elastin, and NAD(P)H emit light in the blue bandpass, while fatty acids and vitamin A emit in the green bandpass. Flavins emit in the yellow bandpass, while pigments and lipofuscins span the visible spectrum. Additionally, porphyrins emit in the red bandpass. Haemoglobin and Mb have known absorbance and emission spectra that have been studied for decades and share porphyrins as a precursor of their synthesis. In 2000, Garry et al. already showed in a review that the pigments were attenuated in the heart of myoglobin knock-out mice ^17^. However, to the best of our knowledge, no biophysical evidence on the involvement of Mb in the endogenous fluorescence of myocardium had been reported previously, and no spectral characterization of the endogenous fluorescence of myocardium has ever been performed. In the current study, we reported that the spatial distribution of Mb in isolated adult cardiomyocytes is distributed similarly to the main endogenous fluorescence in the myocardium. Moreover, we showed that modifications in the redox status of *in vitro* Mb and cleared myocardium fluorescence caused a similar shift in the intensity of the endogenous fluorescence spectrum within the red wavelength bandpass. These collective findings strongly support the conclusion that, in the absence of hemoglobin, Mb is the main contributor to the endogenous fluorescence emission of the myocardium in the red bandpass (>600nm).

### Meaning of Mb oxidation in heart

Auto-oxidation of myoglobin has been extensively reported (refer to ^40^ for review) as oxidation of Mb by different ROS species, including H_2_O_2_, as well ^41^. Interestingly, in 1976, Gotoh and Shikama reported that autoxidation of OxyMb led to a co-oxidation mechanism that generates additional superoxide anion ^42^. Thus, this mechanism could have a deleterious effect on the onset of reperfusion in MI. However, CarbMb has also been reported to exert a nitric reductase activity. Reducing nitrites by CarbMb leads to the generation of MetMb + nitric oxide (NO•), which could, in turn, inhibit mitochondrial respiration and thus induce a protective effect against reperfusion injury ex vivo ^43^.

Furthermore, this mechanism protects the myocardium against oxidation ^44^. Altogether, these results established that MetMb level represents an equilibrium between the oxidation of OxyMb and CarbMb by ROS, auto-oxidation of OxyMb, and the nitrite reductase activity of CarbMb. Finally, its reductase activity can reduce the MetMb level, which converts it back to CarbMb or oxidizes it into ferryl myoglobin (ferrylMb). The latter can be reduced back to MetMb after peroxidation of unsaturated lipids ^40^. The optical absorbance of FerrylMb is similar to that of MetMb, and it is challenging to discriminate between them. However, in the current study, we showed that in excess H_2_O_2_, the emission intensity in the red bandpass is lower than that of a solution enriched with MetMb (**Supplementary Figure S2B**). Depending on the fluorescence spectrum of ferrylMb, the latter result could be explained by a transformation of MetMb in ferrylMb or by the destruction of both MetMb and ferrylMb. Overall, in spectral fluorescence imaging, oxidized MB forms: MetMb and ferrylMb can be discriminated by their intensity in the red bandpass, while CarbMb, and OxyMb are confounded in the absence of spectral analysis in the range 550-620nm.

Previous studies have extensively reported that a burst of ROS production occurs on the onset of reperfusion, which results in significant oxidation of the myocardium during the early phase of reperfusion ^44^. Using oxidation reduction imaging (ORI), the present study estimated that MetMb level increased at 15min post-reperfusion, consistent with the ROS production surge occurring early during reperfusion. Nevertheless, the peak level of MetMb was reached after 3 hours of reperfusion and returned to its basal level after 24 hours. It is unlikely that several minutes of intense ROS production could sustain the oxidation increase until 3h post-reperfusion in the presence of antioxidant enzymes. Noteworthily, the transient dynamics of MetMb level correlated with the changes observed in plasma Troponin I levels, which peaked at 3 hours after reperfusion. Troponin I release into the bloodstream is believed to originate directly from cellular injury/death. As a result, we propose that the transient dynamics of *in situ* MetMb level could be associated with the immediate boost of ROS production by mitochondria on the onset of reperfusion, followed by ROS production from cellular death or inflammatory cells within the initial hours after reperfusion. This transient rise in MetMb was entirely averted when ischemic post-conditioning was performed, confirming that the severity of the infarct is associated with myoglobin’s oxidation.

A part of the MetMb signal may have been related to the nitrite reductase activity induced by NO synthesis by endothelial cells. However, in infarcted hearts exposed to cardioprotection by ischemic post-conditioning, where NO release is expected, the fluorescent signal of MetMb was drastically reduced. We thus inferred that the nitrite reductase activity of myoglobin was not involved in the formation of MetMb during MI.

Interestingly, the oxidized volume, as determined by the increased fluorescence emission in the red bandpass, increased during ischemia and doubled between 15min and 3h post-reperfusion, remaining stable until 24h post-reperfusion. This suggests that the area subjected to a net increase in oxidation begins during ischemia, grows during reperfusion until 3h, and then stabilizes without regressing. This dynamic behavior is expected for the development of the infarcts: initial stress during oxygen and nutriment deprivation, followed by the progressive development of reperfusion injury and stabilization of the necrotic and inflammatory volume. The correlation between MRI and ORI images supports that both techniques measure two correlated mechanisms, namely the diffusion of Gadolinium-DOTA and the MetMb level, and ORI provides an estimate of the infarct size. By combining both the measures of the oxidation intensity and volume, it can be observed that, although the mean intensity of oxidation decreases 24h after reperfusion, there remains a substantial heterogeneity with dense clusters of intense endogenous fluorescence in the red bandpass (as seen in Figure. 4C). These areas may reflect the necrotic regions with dying cardiomyocytes within the heart.

### Myoglobin: a promising imaging biomarker candidate to estimate the oxidation-reduction level in the heart?

The significant advantage of ORI lies in the early diagnosis of the severity of an infarct at the onset of reperfusion. However, the significant limitations include the requirement for tissue sampling and clearing and the penetration of fluorescence light into deeper tissue layers. Despite these drawbacks, there have been advances in cardiovascular molecular imaging, including the use of optical tomography for MI diagnosis ^22^. However, conceptual and technical breakthroughs are required to identify fluorescent targets and improve in-depth-imaging. In the present study, we propose that MetMb is a suitable candidate imaging biomarker to measure the intensity of reperfusion injury within the first hours after reperfusion and could be used to quantify the cardioprotective effect of treatments. Bypassing the current optical limitations is required to translate this seminal work to in vivo animal studies and, eventually, patients. Promising hints for achieving this breakthrough come from photo-acoustics. Lin et al. demonstrated the ability to discriminate between OxyMb and CarbMb in vivo ^45^, suggesting that detection and quantification of MetMb could be achieved using this technique.

### Conclusion

In this study, we developed a novel oxidation/reduction imaging method that quantifies oxidation intensity and oxidized volume during the very early stages of reperfusion following MI. Our findings suggest that using this method can provide an estimate of infarct size comparable to that obtained through MRI. This study may pave the way for developing a novel imaging method for MI diagnosis and follow-up of cardioprotective treatments.

## AUTHOR CONTRIBUTIONS

Sally Badawi, Clémence Leboullenger, Matthieu Chourout, Yves Gouriou, Alexandre Paccalet, Chris Berd, Bruno Pillot, Lionel Augeul, Radu Bolbos, Antonino Bongiovani, Tardivel Meryem, Emmanuelle Canet-Soulas, Claire Crola Da Silva and Gabriel Bidaux conceived and performed experiments.

Sally Badawi, Clémence Leboullenger, Matthieu Chourout, Emmanuelle Canet-Soulas, Claire Crola Da Silva and Gabriel Bidaux wrote the manuscript.

Michel Ovize, Mazen Kurdi, Claire Crola Da Silva and Gabriel Bidaux supervised the study. Nathan Mewton, Thomas Bochaton, Michel Ovize and Mazen Kurdi provided expertise and feedbacks.

Nathan Mewton, Thomas Bochaton, Michel Ovize, Mazen Kurdi and Emmanuelle Canet-Soulas reviewed the manuscript.

## ACKNOWLEDGEMENTS

We would like to thank Mrs Karine Labour and Logos Biosystems for their help and support on improving the clarification of mouse hearts with X-clarity. The authors thank Mr Chris Berd for his experimental help.

## SOURCES OF FUNDING

This work was supported by: IHU OPeRa (ANR-10-IBHU-004) program “Investissements d’Avenir” operated by the French National Research Agency (ANR), RHU MARVELOUS (ANR-16-RHUS-0009) of Université de Lyon, program “Investissements d’Avenir” operated by the French National Research Agency (ANR), INSERM and UCBL annual funding of the CarMeN Laboratory.

## DISCLOSURES

None.

## Notes

### Competing Interest Statement

The authors have declared no competing interest.

## REFERENCES

1. Buja LM. Myocardial ischemia and reperfusion injury. Cardiovasc Pathol Off J Soc Cardiovasc Pathol 2005;14:170–175. doi:10.1016/j.carpath.2005.03.006.

2. Braunwald E, Kloner RA. Myocardial reperfusion: a double-edged sword? J Clin Invest 1985;76:1713–1719.

3. Yellon DM, Hausenloy DJ. Myocardial Reperfusion Injury. N Engl J Med 2007;357:1121–1135. doi:10.1056/NEJMra071667.

4. Reimer KA, Lowe JE, Rasmussen MM, Jennings RB. The wavefront phenomenon of ischemic cell death. 1. Myocardial infarct size vs duration of coronary occlusion in dogs. Circulation 1977;56:786–794. doi:10.1161/01.CIR.56.5.786.

5. Tavares AMV, da Rosa Araujo AS, Llesuy S, Khaper N, Rohde LE, Clausell N, et al. Early loss of cardiac function in acute myocardial infarction is associated with redox imbalance. Exp Clin Cardiol 2012;17:263–267.

6. Garry DJ, Kanatous SB, Mammen PPA. Emerging Roles for Myoglobin in the Heart. Trends Cardiovasc Med 2003;13:111–116. doi:10.1016/S1050-1738(02)00256-6.

7. Millikan GA. MUSCLE HEMOGLOBIN. Physiol Rev 1939. doi:10.1152/physrev.1939.19.4.503.

8. Hendgen-Cotta UB, Kelm M, Rassaf T. A highlight of myoglobin diversity: the nitrite reductase activity during myocardial ischemia-reperfusion. Nitric Oxide Biol Chem 2010;22:75–82. doi:10.1016/j.niox.2009.10.003.

9. Rassaf T, Flögel U, Drexhage C, Hendgen-Cotta U, Kelm M, Schrader J. Nitrite reductase function of deoxymyoglobin: oxygen sensor and regulator of cardiac energetics and function. Circ Res 2007;100:1749–1754. doi:10.1161/CIRCRESAHA.107.152488.

10. Bertelsen G, Skibsted LH. Photooxidation of oxymyoglobin. Wavelength dependence of quantum yields in relation to light discoloration of meat. Meat Sci 1987;19:243–251. doi:10.1016/0309-1740(87)90070-2.

11. Millar SJ, Moss BW, Stevenson MH. Some observations on the absorption spectra of various myoglobin derivatives found in meat. Meat Sci 1996;42:277–288. doi:10.1016/0309-1740(94)00045-X.

12. Anderson AB, Robertson CR. Absorption spectra indicate conformational alteration of myoglobin adsorbed on polydimethylsiloxane. Biophys J 1995;68:2091–2097.

13. Bowen WJ. The absorption spectra and extinction coefficients of myoglobin. J Biol Chem 1949;179:235–245.

14. Roostalu U, Thisted L, Skytte JL, Salinas CG, Pedersen PJ, Hecksher-Sørensen J, et al. Effect of captopril on post-infarction remodelling visualized by light sheet microscopy and echocardiography. Sci Rep 2021;11:5241. doi:10.1038/s41598-021-84812-7.

15. Xu J, Ma Y, Yu T, Zhu D. Quantitative assessment of optical clearing methods in various intact mouse organs. J Biophotonics 2019;12:e201800134. doi:10.1002/jbio.201800134.

16. Des Tombe AL, Van Beek-Harmsen BJ, Lee-De Groot MBE, Van Der Laarse WJ. Calibrated histochemistry applied to oxygen supply and demand in hypertrophied rat myocardium. Microsc Res Tech 2002;58:412–420. doi:10.1002/jemt.10153.

17. Garry DJ, Meeson A, Yan Z, Williams RS. Life without myoglobin. Cell Mol Life Sci CMLS 2000;57:896–898. doi:10.1007/PL00000732.

18. Mukai K, Rosai J, Hallaway BE. Localization of myoglobin in normal and neoplastic human skeletal muscle cells using an immunoperoxidase method. Am J Surg Pathol 1979;3:373–376. doi:10.1097/00000478-197908000-00008.

19. Kamga C, Krishnamurthy S, Shiva S. Myoglobin and Mitochondria: A relationship bound by Oxygen and Nitric Oxide. Nitric Oxide 2012;26:251–258. doi:10.1016/j.niox.2012.03.005.

20. Vinegoni C, Razansky D, Figueiredo J-L, Nahrendorf M, Ntziachristos V, Weissleder R. Normalized Born ratio for fluorescence optical projection tomography. Opt Lett 2009;34:319–321. doi:10.1364/ol.34.000319.

21. Zhao X, Wu J, Gray CD, McGregor K, Rossi AG, Morrison H, et al. Optical projection tomography permits efficient assessment of infarct volume in the murine heart postmyocardial infarction. Am J Physiol-Heart Circ Physiol 2015;309:H702–H710. doi:10.1152/ajpheart.00233.2015.

22. Kramer CM, Sinusas AJ, Sosnovik DE, French BA, Bengel FM. Multimodality imaging of myocardial injury and remodeling. J Nucl Med Off Publ Soc Nucl Med 2010;51 Suppl 1:107S–121S. doi:10.2967/jnumed.109.068221.

23. Leisey JR, Scott DA, Grotyohann LW, Scaduto RC. Quantitation of myoglobin saturation in the perfused heart using myoglobin as an optical inner filter. Am J Physiol 1994;267:H645–653. doi:10.1152/ajpheart.1994.267.2.H645.

24. Femnou AN, Kuzmiak-Glancy S, Covian R, Giles AV, Kay MW, Balaban RS. Intracardiac light catheter for rapid scanning transmural absorbance spectroscopy of perfused myocardium: measurement of myoglobin oxygenation and mitochondria redox state. Am J Physiol - Heart Circ Physiol 2017;313:H1199–H1208. doi:10.1152/ajpheart.00306.2017.

25. Nehrhoff I, Ripoll J, Samaniego R, Desco M, Gómez-Gaviro MV. Looking inside the heart: a see-through view of the vascular tree. Biomed Opt Express 2017;8:3110–3118. doi:10.1364/BOE.8.003110.

26. Smith RM, Matiukas A, Zemlin CW, Pertsov AM. Nondestructive optical determination of fiber organization in intact myocardial wall. Microsc Res Tech 2008;71:510–516. doi:10.1002/jemt.20579.

27. Lee S-E, Nguyen C, Yoon J, Chang H-J, Kim S, Kim CH, et al. Three-dimensional Cardiomyocytes Structure Revealed By Diffusion Tensor Imaging and Its Validation Using a Tissue-Clearing Technique. Sci Rep 2018;8. doi:10.1038/s41598-018-24622-6.

28. Wang Z, Zhang J, Fan G, Zhao H, Wang X, Zhang J, et al. Imaging transparent intact cardiac tissue with single-cell resolution. Biomed Opt Express 2018;9:423. doi:10.1364/BOE.9.000423.

29. Merz SF, Korste S, Bornemann L, Michel L, Stock P, Squire A, et al. Contemporaneous 3D characterization of acute and chronic myocardial I/R injury and response. Nat Commun 2019;10:2312. doi:10.1038/s41467-019-10338-2.

30. Harris SM, Harvey EJ, Hughes TR, Ramji DP. The interferon-γ-mediated inhibition of lipoprotein lipase gene transcription in macrophages involves casein kinase 2- and phosphoinositide- 3-kinase-mediated regulation of transcription factors Sp1 and Sp3. Cell Signal 2008;20:2296–2301. doi:10.1016/j.cellsig.2008.08.016.

31. Whitburn KD. The interaction of oxymyoglobin with hydrogen peroxide: the formation of ferrylmyoglobin at moderate excesses of hydrogen peroxide. Arch Biochem Biophys 1987;253:419–430. doi:10.1016/0003-9861(87)90195-0.

32. Yamashita T, Abe S, Arima S, Nomoto K, Miyata M, Maruyama I, et al. Myocardial infarct size can be estimated from serial plasma myoglobin measurements within 4 hours of reperfusion. Circulation 1993;87:1840–1849. doi:10.1161/01.cir.87.6.1840.

33. Stone MJ, Waterman MR, Poliner LR, Templeton GH, Buja LM, Willerson JT. Myoglobinemia is an early and quantitative index of acute myocardial infarction. Angiology 1978;29:386–392. doi:10.1177/000331977802900506.

34. Hachey BJ, Kontos MC, Newby LK, Christenson RH, Peacock WF, Brewer KC, et al. Trends in Use of Biomarker Protocols for the Evaluation of Possible Myocardial Infarction. J Am Heart Assoc 2017;6:e005852. doi:10.1161/JAHA.117.005852.

35. Kottwitz J, Bruno KA, Berg J, Salomon GR, Fairweather D, Elhassan M, et al. Myoglobin for Detection of High-Risk Patients with Acute Myocarditis. J Cardiovasc Transl Res 2020;13:853–863. doi:10.1007/s12265-020-09957-8.

36. Ti Y, Chen P, Lin W-C. In vivo characterization of myocardial infarction using fluorescence and diffuse reflectance spectroscopy. J Biomed Opt 2010;15:037009. doi:10.1117/1.3442505.

37. Takakura H, Masuda K, Hashimoto T, Iwase S, Jue T. Quantification of myoglobin deoxygenation and intracellular partial pressure of O2 during muscle contraction during haemoglobin-free medium perfusion. Exp Physiol 2010;95:630–640. doi:10.1113/expphysiol.2009.050344.

38. Zhao X, Wu J, Gray CD, McGregor K, Rossi AG, Morrison H, et al. Optical projection tomography permits efficient assessment of infarct volume in the murine heart postmyocardial infarction. Am J Physiol Heart Circ Physiol 2015;309:H702–710. doi:10.1152/ajpheart.00233.2015.

39. Croce AC, Bottiroli G. Autofluorescence spectroscopy and imaging: a tool for biomedical research and diagnosis. Eur J Histochem EJH 2014;58:2461. doi:10.4081/ejh.2014.2461.

40. Richards MP. Redox reactions of myoglobin. Antioxid Redox Signal 2013;18:2342–2351. doi:10.1089/ars.2012.4887.

41. Yusa K, Shikama K. Oxidation of oxymyoglobin to metmyoglobin with hydrogen peroxide: involvement of ferryl intermediate. Biochemistry 1987;26:6684–6688. doi:10.1021/bi00395a018.

42. Gotoh T, Shikama K. Generation of the superoxide radical during autoxidation of oxymyoglobin. J Biochem (Tokyo) 1976;80:397–399. doi:10.1093/oxfordjournals.jbchem.a131289.

43. Hendgen-Cotta UB, Merx MW, Shiva S, Schmitz J, Becher S, Klare JP, et al. Nitrite reductase activity of myoglobin regulates respiration and cellular viability in myocardial ischemia-reperfusion injury. Proc Natl Acad Sci U S A 2008;105:10256–10261. doi:10.1073/pnas.0801336105.

44. Flögel U, Gödecke A, Klotz L-O, Schrader J. Role of myoglobin in the antioxidant defense of the heart. FASEB J Off Publ Fed Am Soc Exp Biol 2004;18:1156–1158. doi:10.1096/fj.03-1382fje.

45. Lin L, Yao J, Li L, Wang LV. In vivo photoacoustic tomography of myoglobin oxygen saturation. J Biomed Opt 2016;21:61002. doi:10.1117/1.JBO.21.6.061002.

